# The human *FLT1* regulatory element directs vascular expression and modulates angiogenesis pathways *in vitro* and *in vivo*

**DOI:** 10.1101/2021.03.03.433738

**Authors:** Julian Stolper, Holly K. Voges, Michael See, Neda Rahmani Mehdiabadi, Gulrez Chahal, Mark Drvodelic, Michael Eichenlaub, Tanya Labonne, Benjamin G. Schultz, Alejandro Hidalgo, Lazaro Centanin, Jochen Wittbrodt, Enzo R. Porrello, David A. Elliott, Mirana Ramialison

## Abstract

There is growing evidence that mutations in non-coding *cis*-regulatory elements (CREs) disrupt proper development. However, little is known about human CREs that are crucial for cardiovascular development. To address this, we bioinformatically identified cardiovascular CREs based on the occupancy of the CRE by the homeodomain protein NKX2-5 and cardiac chromatin histone modifications. This search defined a highly conserved CRE within the *FLT1* locus termed *enFLT1*. We show that the human *enFLT1* is an enhancer capable of driving reporter transgene expression *in vivo* throughout the developing cardiovascular system of medaka. Deletion of the human *enFLT1* enhancer (*ΔenFLT1*) triggered molecular perturbations in extracellular matrix organisation and blood vessel morphogenesis *in vitro* in endothelial cells derived from human embryonic stem cells and vascular defects *in vivo* in medaka. These findings highlight the crucial role of the human *FLT1* enhancer and its function as a regulator and buffer of transcriptional regulation in cardiovascular development.

## Introduction

Disruption of heart and major blood vessel formation during development results in congenital heart defects at birth and are a major factor underlying child mortality and morbidity (1). The development of the heart is strictly regulated by a tight network of genetic components (2) which when disrupted perturb normal heart development leading to disease. Genome wide association studies (GWAS) have been used to identify genes associated with CHD and begin to dissect the complex genetic architecture underlying heart development (3,4). Nevertheless, an established problem is definitively assigning pathogenicity to a given variant or single nucleotide polymorphism (SNP). Strikingly, the majority of SNPs associated with CHD are found in non-coding regions in the genome, highlighting the importance of *cis*-regulatory elements in developmental processes and disease (5,6). These regulatory elements (REs), such as enhancers, promote gene expression in a spatial temporal manner through the coordinated binding of specific transcription factors (TFs). However, since SNPs in such sequences can lead to disruptions in TF binding motifs and therefore have no impact on the protein sequence directly, it is challenging to pinpoint the how sequence alterations result in cardiac and blood vessel defects. Aberrations in regulatory elements can lead to the perturbation of a gene regulatory network which, in turn, causes genes to be over- or underexpressed, even for multiple targets at once. This ultimately results in enhanceropathies, a group of diseases caused by mutations in regulatory elements (7).

There is now mounting evidence that disrupted cardiovascular regulatory elements can impair heart development leading to disease (8–11). A prerequisite for functional studies to understand the effect of non-coding SNPs, is the accurate identification of the regulatory regions in the human genome that are important for cardiac development and disease. Accessible datasets of the human genome, regulatory element associated chromatin marks (12,13), TF analyses (14) and chromatin capture experiments (15–17) provide valuable resources to define the human gene regulatory network in the heart. However, key challenges to identify non-coding elements relevant for disease remain. The search space is still large, for example, the current registry of human *cis*-regulatory elements in the Encode data set is comprised of 926,535 entries (18). Furthermore, it is crucial to have functional validation methods to determine both the sufficiency and necessity of a given human regulatory element for normal development (19).

In this study, we developed a bioinformatic pipeline to identify cardiac enhancers that are involved in development and disease with a particular focus on the highly conserved cardiac TF NKX2-5. NKX2-5 is essential for heart formation and homeostasis and is crucial for the development of heart muscle cells (14,20,21). Since mutations in NKX2-5 can lead to CHD (22), we reasoned that variants in NKX2-5 target enhancers may also impair cardiac development. By using datasets of histone modifications, evolutionary conservation and functional enrichment, we have identified a human enhancer of the *Fms Related Receptor Tyrosine Kinase 1* gene (*FLT1* also known as *Vascular growth factor receptor1* (*VEGFR1*)) termed *enFLT1*. Dysregulation of the *FLT1* genes leads to vascular abnormalities and cardiovascular phenotypes in fish, mice and humans (23–25). Despite extensive study of *FLT1* function, its regulatory network of enhancers has not been completely defined (26–28). How perturbations of these elements may alter the transcriptional regulation of *FLT1* has yet to be determined. Here we demonstrate that the human *enFLT1* enhancer was able to drive gene expression in cardiovascular tissues *in vivo* in medaka (*Oryzias latipes*). The deletion of the enhancer element in endothelial cells derived from human embryonic stem cells revealed a molecular disruption which overlaps with *FLT1* gene loss-of-function. In addition, medaka *enFLT1* deletion resulted in impaired cardiovascular development *in vivo*. Thus, here were have defined a highly conserved *FLT1* enhancer and provide evidence that this enhancer plays an evolutionarily important role in the development of the cardiovascular system through the modulation of *FLT1* downstream pathways essential for blood vessel morphogenesis.

## Results

### Identification of *cis*-regulatory elements relevant for cardiac development and disease

In order to identify human *cis*-regulatory elements that are involved in cardiovascular development, we developed a bioinformatic pipeline to filter for sequences that were directly bound by NKX2-5, a TF essential for heart development (Fig 1a). We therefore made use of a previously generated dataset of NKX2-5 genomic targets identified in human pluripotent derived cardiomyocytes by chromatin immunoprecipitation sequencing (ChIP-seq) (14). From all of the ChIP-seq experiments, 20,879 regions were identified to be directly bound by NKX2-5. Since heart development is a highly conserved process, REs deeply embedded in such essential processes are under positive selective pressure compared to non-functional non-coding sequences to maintain correct activity (12). We therefore filtered these regions for high sequence conservation and obtained 62 sequences which were ultra-conserved across 100 vertebrate species, from fish to human. Furthermore, in order to filter for sequences that were shown to be active REs, we used publicly available datasets of histone modification marks as a measure to obtain active cardiac enhancers(13). We identified 38 regions that showed histone marks for active enhancers H3K4 monomethylation (H3K4me1) and H3K27 acetylation (H3K27ac). We further filtered these regions for those that could be associated with genes known to play a role in heart development. We identified 7 CREs associated with highly relevant genes expressed in the heart and deeply embedded in the genetic networks controlling heart development (Table S1). To further understand the mechanism of regulatory elements involved in cardiovascular development and to illustrate the evidence supporting our hypothesis, we set out to investigate an enhancer element located in the intron 10 of the gene *FLT1* (Fig. 1b).

**Figure 1:**
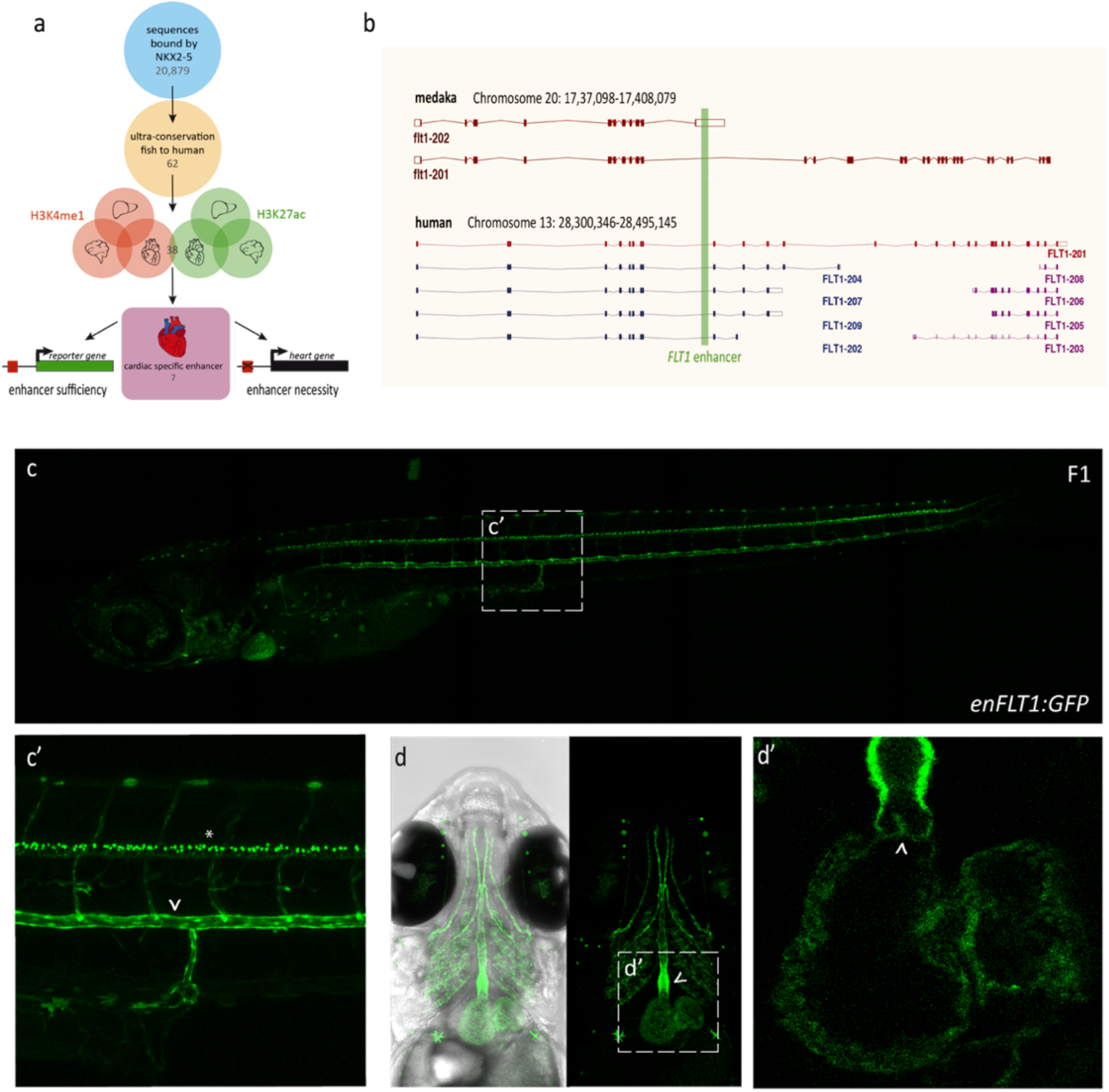
Identification and validation of cardiac specific *cis*-regulatory elements. a) Bioinformatic pipeline to identify cardiac regulatory elements (RE) based on NKX2-5 binding, sequence conservation and active histone marks (H3K4me1 and H3K27ac). Putative RE were tested for the ability to drive reporter gene expression and the necessity for endogenous *FLT1* expression. b) Overview of the *FLT1* locus and its isoforms in humans and medaka. The RE used for *in vivo* transgenesis is displayed in green. c) *enFLT1* drives stable reporter gene (green fluorescent protein (GFP)) expression in neurons (asterisk) and endothelium such as c’) the dorsal aorta (arrowhead) d) the outflow tract (arrowhead) d’) and valves (arrowhead) in the heart.

### The human enhancer of *FLT*1 is able to drive cardiovascular gene expression *in vivo*

We set out to assess the *in vivo f*unction of the human enhancer sequence *enFLT1*, which has been validated to be bound by many TFs embedded in cardiovascular development (Fig. S1a). In order to determine whether the RE is able to drive GFP reporter gene expression in the heart, we cloned the enhancer in a modified ZED vector (29,30) to perform a transgenesis assay in medaka (Fig. S1b). Already established in the ENCODE project in 2012, medaka has been shown to be the ideal model to study human regulatory elements *in vivo* and is well suited for genetic engineering (12). Of 121 injected embryos, 52 embryos (43%) showed consistent GFP expression in the cardiovascular system (Fig. S1c). All GFP positive fish were raised and subsequently crossed to wildtype fish to obtain a stable transgenic line (Fig. 1c) denoted *enFLT1:GFP*. Characterisation of the *enFLT1:GFP* line revealed consistent expression in cardiac and endothelial tissues such as intersegmental vessels, dorsal aorta (Fig. 1c’), the outflow tract (Fig. 1d), blood vessels in the myocardium, the endocardium and the heart valves (Fig. 1d’). This demonstrates that *enFLT1* consisting of 358 bases is sufficient to drive GFP expression in endothelial and cardiac tissues *in vivo*.

### Enhancer of *FLT1* is essential for pathways involved in blood vessel morphogenesis

To understand the effect of the *FLT1* enhancer on the gene regulatory network and its function in humans, we deleted the RE via CRISPR/Cas9 mediated gene editing (Fig. S2a) in human embryonic stem cells (hESC) (background line: H3, NKX2-5(^eGFP/wt^) (31)) to generate the Δen*FLT1* line, which was subsequently differentiated into endothelial cells. In order to understand the effects of the enhancer deletion on FLT1 and its related pathways, we also generated a gene mutant cell line, Δex1*FLT1* to act as a positive control. To understand the transcriptional consequences of deleting *enFLT1* (Fig. 2a) we performed RNA sequencing on wildtype, *ΔenFLT1* and *Δex1FLT1* endothelial cells derived from hESC after 12 days of culturing. Deletion of the enhancer had only a minor, not significant reduction on *FLT1* mRNA expression (Fig. S2b) despite evidence from chromatin conformation capture data (32) that enFLT1 interacts with the FLT1 promoter in cardiac cells (Fig. S2c). The removal of exon 1 of the gene however resulted in a truncated transcript (Fig. S2d). Nonetheless, when assessing the expression of different *FLT1* isoforms in *ΔenFLT1*, we could not detect a significant decrease in transmembrane bound *FLT1 (tFLT1)* expression (201) nor any difference in expression levels of any of the soluble *FLT1 (sFLT1)* isoforms (204/207), suggesting a more redundant, subtle role of the CRE in *FLT1* regulation. However, deletion of exon 1 of *FLT1* had a significant effect on expression of both, transmembrane and soluble isoforms (Fig. 2b), highlighting *FLT1* dysregulation. Furthermore, we identified 211 differentially expressed genes (DEGs) as a result of deleting the regulatory element compared to cells with the intact wildtype *enFLT1*. The majority of these DEGs (179) overlapped with the 2261 identified genes which were differentially expressed in the human *FLT1* gene mutant, suggesting an essential role of the enhancer in the FLT1 pathway (Fig. 2c). Furthermore, for the first time, we looked at gene ontology (GO) terms of the DEGs in a human *FLT1* gene mutant and found that the main pathways affected were blood vessel development, cell differentiation, cell division, heart development and extra cellular matrix organisation consistent with the molecular function of FLT1 (25,33). The genes shared by gene mutant and enhancer mutant human cell lines were involved in pathways such as organelle fission and extracellular structure organisation and other regulatory processes of extracellular matrix (ECM) composition. Pathways identified to be unique to differentially expressed genes in *ΔenFLT1* were also found to be involved in angiogenesis and blood vessel morphogenesis, suggesting a role of *enFLT1* in alternative pathways in blood vessel development distinct from the FLT1 pathway (Fig. 2d and S2e).

**Figure 2:**
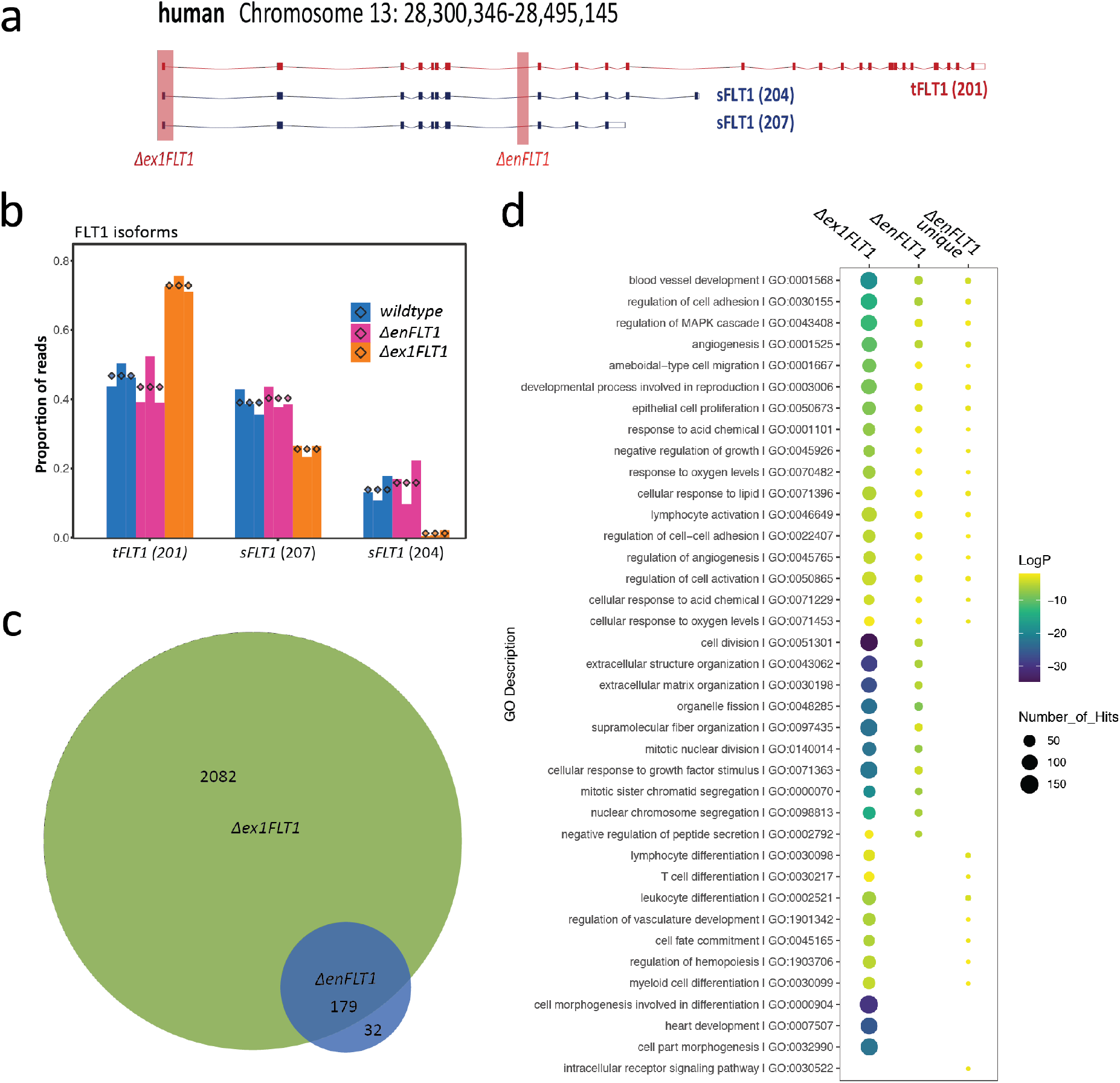
*enFLT1* is implemented in FLT1 related and alternative pathways. a) Overview of the transmembrane bound FLT1 (tFLT1) and soluble FLT1 (sFLT1) isoforms. We generated a gene mutant (*Δex1FLT1*) and enhancer mutant (*ΔenFLT1*) cell line in human embryonic stem cells. b) Differentially expressed isoforms in *wildtype, ΔenFLT1* and *Δex1FLT1* cells. c) Venn diagram of differentially expressed genes (DEGs) (compared to wildtype) in *ΔenFLT1* and *Δex1FLT1*. The majority of DEGs in *ΔenFLT1* overlaps with genes in *Δex1FLT1*. d) Gene ontology of DEGs in Δex1FLT1, ΔenFLT1 and the 32 genes unique to enhancer mutant.

### Deletion of *enFLT1* does not impair angiogenesis *in vitro*

In order to assess the potential of *ΔenFLT1* and *Δex1FLT1* endothelial cells to form tubes *in vitro*, we performed angiogenesis assays (34). Cells were labelled with anti-human CD31 and anti-human CD34 antibodies and imaged over the course of 48 hours (Fig. 3a). While wildtype (*NKX2-5*^*eGFP/wt*^) and *ΔenFLT1* endothelial cells formed tubes within 10h and then stopped and coalesced, we observed a continuation of angiogenesis and proliferation in *Δex1FLT1* cells until 48h (Fig. 3b). To assess difference in tube formation potential across genotypes, we analysed skeleton length, area, junction count and branch count over the course of 48h (Fig. 3c and S3a). *ΔenFLT1* cells showed no significant difference compared to wildtype cells in any of the categories assessed. However, we did see a significant change in area and skeleton length when comparing *Δex1FLT1* with the other cell lines (Fig. 3d), indeed suggesting a disruption of the *FLT1* pathway during angiogenesis consistent with published data on *FLT1* knockdown in HUVECS and *Flt1*^*-/-*^mutant in endothelial cells derived from murine embryonic stem cells (23). This data suggests the enhancer of *FLT1* is not essential for endothelial tube formation *in vitro* and that other regulatory elements act to buffer *FLT1* expression from the loss of *enFLT1*. Nevertheless, given the altered transcriptional profile observed in *ΔenFLT1* endothelial cells it remains possible that this enhancer is required for normal development.

**Figure 3:**
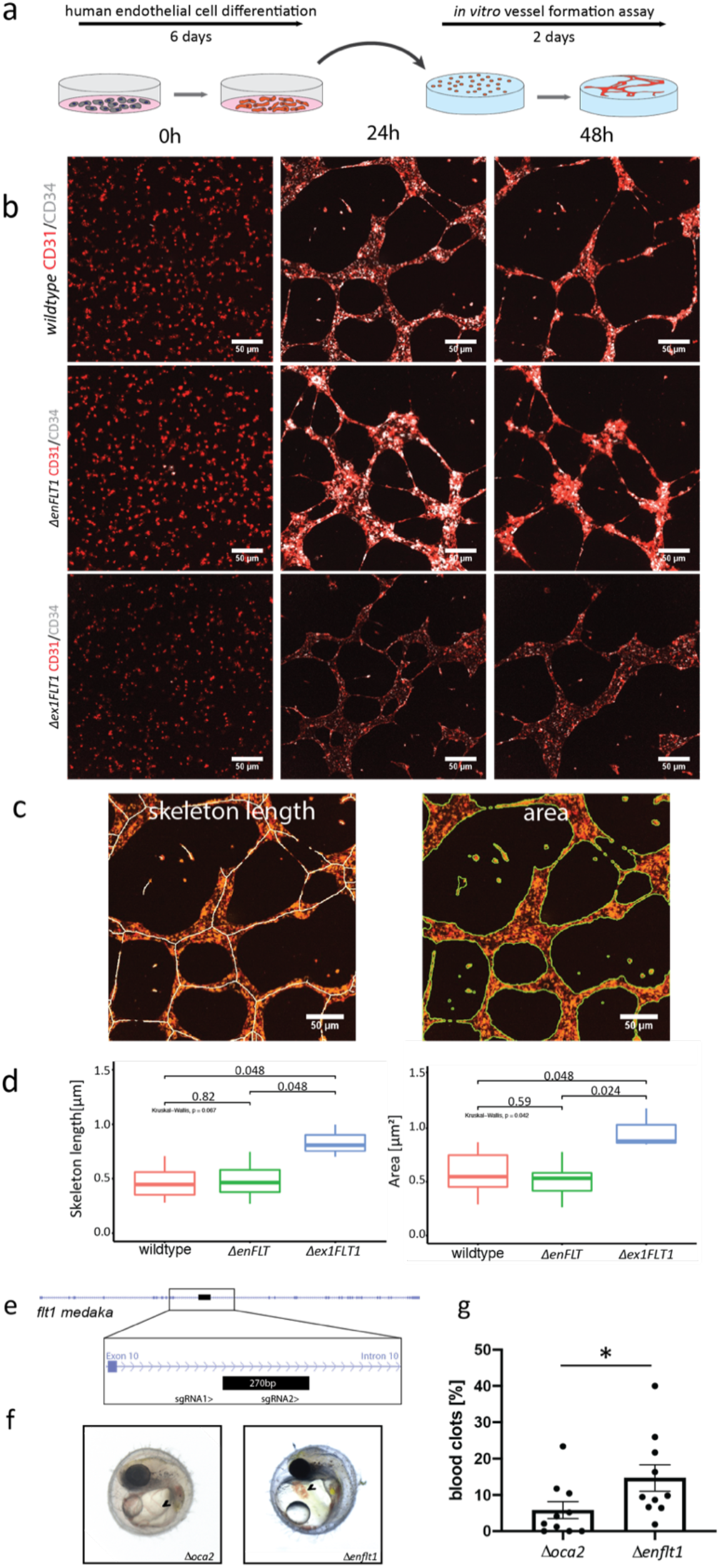
Functional analysis of *enFLT1* during angiogenesis. a) human embryonic stem cells were differentiated within 6 days and subsequently used for angiogenesis assays and imaged for 48 hours. b) Angiogenesis assays of wildtype, *ΔenFLT1* and *Δex1FLT1* cell lines at 0h, 24h, and 48h. All cell lines stayed positive for CD31 (red) and CD34 (grey) for 48h. *Δex1FLT1* vessels maintained stronger connections and collapsed over time forming thicker vessels. c) CellPathfinder analysis of vessels based on skeleton length and area. d) Statistical analysis of normalised skeleton length and area covered over the course of the experiment. *Δex1FLT1* cells displayed longer connections and covered a significant larger area compared to wildtype and *ΔenFLT1* cells. e) *flt1* locus and endogenous *enflt1* in medaka. 270bp were targeted with 2 sgRNAs. f) Deletion of *enflt1* increased occurrence of blot clots (*Δenflt1*) compared to control injections targeting *oca2* (*Δoca2*) g) statistical analysis of blood clot occurrence. A significant difference (p= 0.0215) of blood clot numbers were observed upon deletion of *enflt1* (14.68%) compared to controls (5.81%).

### Deletion of *enFLT1* perturbs cardiovascular development *in vivo*

To determine if loss of the conserved *FLT1* enhancer alters vasculogenesis *in vitro* we deleted the endogenous en*flt1* in medaka and assessed the effect on blood vessel development or morphology. In order to alter the regulatory sequence, which was located in intron 10 of the *tflt1* isoform and the 3’UTR of the *sflt1* isoform in medaka (Fig. 1b), we made use of the CRISPR/Cas9 System to introduce a 270 bp deletion (Fig. 3e). The excision of the endogenous RE fragment in medaka (*enflt1*) was verified by PCR (Fig. S3b) and injected embryos were analysed for defects in heart or blood vessel formation. To rule out an effect of the Cas9 enzyme or the process of microinjection itself on heart and blood vessel formation, controls were injected to target *oca2*, a gene important for pigment formation in the eye and body (35). Interestingly, embryos injected with the sgRNA targeting *enflt1* (Fig. 3f and g) resulted in a significant higher number of blood clots (14.68%) on the yolk in compared to control embryos (5.81%) (p value=0.0215, Mann-Whitney test, one-tailed) suggesting a crucial role of this sequence in angiogenesis and vascularisation *in vivo* in fish.

## Discussion

FLT1 is an important regulator of blood vessel development, cell proliferation, migration, differentiation and cell survival (36). Loss of FLT1 or perturbation of *Flt1* regulation has been implicated in blood vessel defects in human, mouse, and zebrafish (23–25). Given that blood vessels are throughout the body, *FLT1* regulation is likely to be complex incorporating both tissue specific cues and endothelial cell type signals. Indeed, the precise combination of transcriptional regulators that bind at the regulatory elements of the *FLT1* locus are not well understood. However, a few studies have highlighted the importance of *FLT1 cis*-regulatory elements and suggested that disruptions in these non-coding regulators affect gene expression of *FLT1* and its isoforms and can ultimately lead to disease (26,28,37,38).

We aimed to functionally test regulatory elements which were linked to cardiovascular development and disease in humans and identified an enhancer in intron 10 of *FLT1*. This enhancer is well conserved across species suggesting it plays a critical role in modulating *FLT1* expression levels. For example, the endogenous zebrafish *enFlt1* enhancer in combination with the endogenous *flt1* promoter is able to drive transgenic reporter gene expression throughout the developing vasculature *in vivo* (27). Here we demonstrate that the human *enFLT1* was also able to drive stable reporter gene expression in combination with a synthetic minimal promoter (SCP1) (39). In human endothelial cells derived from *ΔFLT1* hESCs levels of *FLT1* were only slightly reduced from wildtype levels. Previous studies have shown that the deletion of *cis*-regulatory elements, despite their capacity to drive gene expression, may only lead to subtle effects on transcription levels or phenotype (40). This occurs due to the fact that genomic regulatory domains often act additively to provide genetic and phenotypic robustness during the development of an organism (9,40–43). Therefore, we hypothesize that other regulatory elements in the locus functionally compensate for *enFLT1* to maintain stable *FLT1* expression. Nevertheless, the transcriptional profile of endothelial cells is perturbed in the *enFLT1* knockout suggesting the enhancer may provide robustness during vessel morphogenesis. Supporting this idea is that the *in vivo* knockout of the endogenous conserved enhancer in medaka led to an increase of blood vessel disruptions on the yolk resulting in blood clot formation, hinting towards an important role of this sequence in vessel morphogenesis in fish (9,40–43).

Using an *in vitro* angiogenesis model we observed vessel collapse and a reduced rate of retraction of vessels in the human *FLT1* gene mutant cell line that was consistent with previous reports (23), however, this phenotype was not recapitulated in *ΔenFLT1* lines. These *in vitro* experiments might be not be the appropriate assay of enhancer function. An *in vivo* mammalian system could provide more insight into determining if the *FLT1* enhancer is integrated into transcriptional networks during organogenesis rather than angiogenesis. For example, the enhancer may have key regulatory roles during cardiac development given the strategy to identify the enhancer used sequences targeted by the cardiac transcription factor NKX2-5. Interestingly we do see an effect of the deletion of the RE in medaka *in vivo*. In fish, the conserved enhancer lies in the 3’UTR of the *sflt1* isoform, which might lead to premature degradation of the mRNA and, therefore, may have a stronger effect on vessel morphogenesis.

Genes differentially expressed in ΔenFLT1 are involved in blood vessel morphogenesis and cardiovascular development via the regulation of the extracellular matrix composition.

Supporting the notion that the *enFLT1* is a critical regulatory element we also observed 33 DEGs, which did not overlap with DEGs in *Δex1FLT1*, that require an intact enhancer. Among these genes, ANGPT1 is involved in angiogenesis and disruptions in the gene have been associated with angioedema (44,45). As an antagonist of angiogenesis, FLT1 has been predicted to be a functional partner of ANGPT1 (46). While the most straightforward explanation is that sub-optimal *FLT1* expression in ΔenFLT1 endothelial cells impairs these pathways, a more speculative hypothesis is that the enhancer acts in *trans* to regulate this subset of genes. However, in order to fully understand the interaction network of the enhancer with other genes, 3D interaction data of the enhancer is required to identify putative physical interaction partners.

In this study we have defined an evolutionarily conserved enhancer element that permits further dissection of the complex regulatory program controlling of *FLT1* expression. This study provides novel cellular and animal models to further study this enhancer element and provides a striking example of the robustness of the transcriptional network the enables the precise expression of *FLT1*. In conclusion, we show that perturbations in the sequence of a regulatory element of *FLT1* compromises the FLT1/VEGF signalling cascade impairing both transcriptional profiles and blood vessel formation. In this context, this work provides a framework for identifying other *FLT1* regulatory sequences that facilitate the complex interplay of spatial and temporal cues provided to the cells of the vasculature throughout the body.

## Methods

### Bioinformatic mining and enhancer prediction

Sequences bound by NKX2-5 were retrieved from Anderson et al (14), filtered for ultra-conserved sequences via MultiZ alignments (47) and intersected with histone modification marks datasets the Human Roadmap Epigenome Project (48). The GREAT tool (49) was used to link enhancer candidates to target genes based on proximity rules.

### Fish maintenance and ethics

Fish lines were maintained under standard recirculating aquaculture conditions. Day-night cycles were set to 14 h of light and 10h of darkness. The whole fish facility is under the supervision of the local representative of the animal welfare agency and all experiments on Medaka (*Oryzias latipes*) were performed according to European Union animal welfare guidelines and national animal welfare standards in Germany (Tierschutzgesetz §11, Abs. 1, Nr.1, husbandry permit number AZ35-9185.64/BH Wittbrodt, line generation permit number AZ 35-9185.81/G-145-15). The wildtype strain Cab was used in this study.

### Enhancer assay and generation of transgenic lines

The reporter line *enFLT1:GFP* was generated by injection of 5ng/µl donor DNA and 10 ng/µl Tol2 transposase mRNA into one cell stage medaka embryos. For transgenesis using the Meganuclease SceI, the injection mix consisted of 0.5x Yamamoto buffer, 0.5x I-SceI buffer, 0.3 U/*μ*l I-SceI enzyme and 10-20n/µl of the donor constructs. The injected plasmid was a modified zebrafish enhancer detection plasmid (29,30) containing a SCP1 promoter (39). The enhancer element was amplified using these primers: trans_enflt1_for: TTAGGGGGAGGGGAATGTGC; trans_enflt1_rev: CCTCCCTGCCATTGTACTTGG.

### Imaging *in vivo*

Medaka hatchlings (sedated with tricaine when alive) were mounted in 1% low melting agarose on glass bottom MatTek dishes. High-resolution imaging was carried out using confocal laser scanning microscopes (Leica TCS SPE or Leica TCS SP8). Analysis and processing were performed using the ImageJ software.

### Gene editing *in vivo*

Suitable sgRNAs with low predicted off targets were designed using the CRISPR/Cas9 target online predictor (CCTop) (50). Cas9 mRNA was transcribed from JDS246 by mMessage mMachine Sp6 Transcription Kit (Thermo Fisher) and sgRNA were cloned into DR274 (Addgene #42250) (51). DR274 was linearized using the restriction enzyme DraI and subsequently transcribed using MEGAscript T7 transcription Kit (Thermo Fisher). RNA purification was performed using the RNeasy Mini kit (Qiagen). One-cell medaka embryos were injected with 150 ng/µl Cas9 mRNA and 15 ng/µl per sgRNA used. Δenflt1_sgRNA1: CCAGACCCAACAGTGGACCC; Δenflt1_sgRNA2: GGGCTTGAGAGGTATGTGCT; Δoca2_sgRNA1: TTGCAGGAATCATTCTGTGT; Δoca2_sgRNA2: GATCCAAGTGGAGCAGACTG (35).

### Gene editing *in vitro*

sgRNA oligos were cloned into px458 (Addgene #48138) and subsequently transfected via electroporation using 100µl Neon^®^ Tips. Human embryonic stem cells were electroporated at 1050V for 30ms with 2 pulses. Cells transiently expressing GFP were single cell sorted, colonies were grown for 2 weeks and screened via PCR. ΔenFLT1_sgRNA1: TAAGGGCACAAGCCCTAGTA; ΔenFLT1_sgRNA2: ACCTGAAACAACTTAATTT; Δex1FLT1_sgRNA1: TAGTTGCAGCGGGCACGCTT; Δex1FLT1_sgRNA2: TTATAAATCGCCCCCGCCCT.

### Cell culture and endothelial cell differentiation

HESCs (background line: H3, NKX2-5(^eGFP^/^wt^)) were cultured on feeders and passaged as previously described (31). Endothelial cell differentiation was induced by using an adapted cardiomyocyte protocol (14). 2.2 × 10^6^ cells were seeded per well on 6-well plates coated with Geltrex (Life Technologies). On day 0, the standard cardiac differentiation medium contained 12µM CHIR99021, 80ng/ml Activin A and 50µg/ml Ascorbic Acid. For day 3 and day 5, the standard media was supplemented with 5µM IWR-1, 50µg/ml Ascorbic Acid, 30ng/ml VEGF (PeproTech) and 50ng/ml SCF (PeproTech). At day 6 the cells were harvested and processed for flow cytometry sorting. The suspension was stained for the endothelial specific cell surface markers CD31 and CD34 to sort for endothelial progenitors. The antibodies anti-human CD31-APC (BioLegend) and anti-human CD34-PE/Cy7 (BioLegend) were used in a 1:400 ratio. Endothelial cells were maintained in endothelial growth medium, EGM2-2MV Bulletkit (Lonza).

### Angiogenesis assay

Vessel formation assays or angiogenesis assays were performed as previously published (34). In brief, 40µl of Geltrex™ were added to each well of a 96 well glass bottom plates (Corning). The plates were kept on ice while pipetting, subsequently centrifuged and left to set in the incubator at 37°C for 30min. Endothelial cells were harvested and resuspended in complete growth factor EGM-2V media supplemented with anti-human CD31-APC and anti-human CD34-PE/Cy7 antibodies (1:400). 15,000 cells were plated per well.

### Imaging and Image analysis

Confocal images were acquired with Yokagawa CellVoyager CV8000 high-throughput discovery system under 37°C and 5% CO2. Maximum intensity projection (MIP) images were constructed from 15µm z slices (300µm total z distance), captured every 40 minutes for 48 hours. Images were analysed in CellPathfinder software based on CD34-APC intensity. Analysis algorithm was defined using skeleton function to generate the parameters vessel area, vessel length, branch count, and junction count. The Kruskal-Wallis test was used to evaluate differences in medians among three cell lines. If the Kruskal-Wallis test was significant, the Wilcoxon Rank Sum Test was used to assess the level of difference significance on a variable between two cell lines. The statistical analysis was performed in the R statistical programming language.

### RNA sequencing and analysis

Three biological replicates with each three technical replicates (per cell line) of endothelial cells were harvested 12 days after differentiation initiation. RNA was extracted using the Direct-Zol RNA Miniprep Kit (Zymo research). Paired-end mRNA sequencing was performed at the

Victorian Clinical Genetics Services (VCGS) using the NovaSeq 6000 System (Illumina) with a 2 x 150 bp read length. The fastq files were processed using the RNAsik pipeline (52). The STAR aligner (53) was used to align reads to the GRCh38 Assembly. Aligned reads were assigned to features from the GRCh38 EnsEMBL Annotation (54) using the featureCounts program from RsubRead (55). Degust (56) was used to perform and visualise differential expression analysis. Firstly, the first dimension of unwanted variation were removed from counts using RUVr routine from the RUVSeq R package (57). Next, TMM normalisation and the quasi-likelihood test was performed using EdgeR’s (58) standard workflow.

Salmon (59) was used to quantify transcript isoforms abundance, and the differential abundance of FLT1 was tested and visualised using the DRIMSeq R Package (60).

## Declarations

### Ethics approval and consent to participate

No ethics approval and consent required for this study.

### Consent for publication

All authors provide consent for publication.

## Authors’ contributions

MR designed the study with input from DE and ME. JS performed *in vivo* and *in vitro* experiments with input from TL, LC, JW, EP, DE and MR. JS and HV performed imaging and data analysis with input from AH. MR, ME, MD, MS, JS, NRM and BS performed bioinformatics and statistical analysis. JS, MR and DE wrote the manuscript with input from all authors. All authors reviewed and approved the manuscript.

## Availability of data and materials

The RNA-seq dataset is accessible through NCBI Gene Expression Omnibus, GEO160873.

## Acknowledgments

We thank Jeannette Hallab, Karen Gross, Kathy Karavendzas, Francesca Bolk, Markus Tondl, Ling Qian and Sebastian-Alexander Stamatis for support in laboratory work and advice. Choon Boon (Evangelyn) Sim, Ali Seleit, James McNamara and Christine Wells for scientific discussions and guidance. We are grateful to Jose Arturo Gutierrez-Triana for providing the transgenesis vector. We would also like to thank the Victorian Clinical Genetics Services (VCGS) for providing the RNA-seq service and the MCRI FACS facility for technical assistance. Furthermore, we would like to thank all members of the Centanin, Elliott, Porrello, Wittbrodt and Ramialison laboratory members for active discussions and feedback. The Australian Regenerative Medicine Institute is supported by grants from the State Government of Victoria and the Australian Government.

## Supplementary data

**Figure S1:**
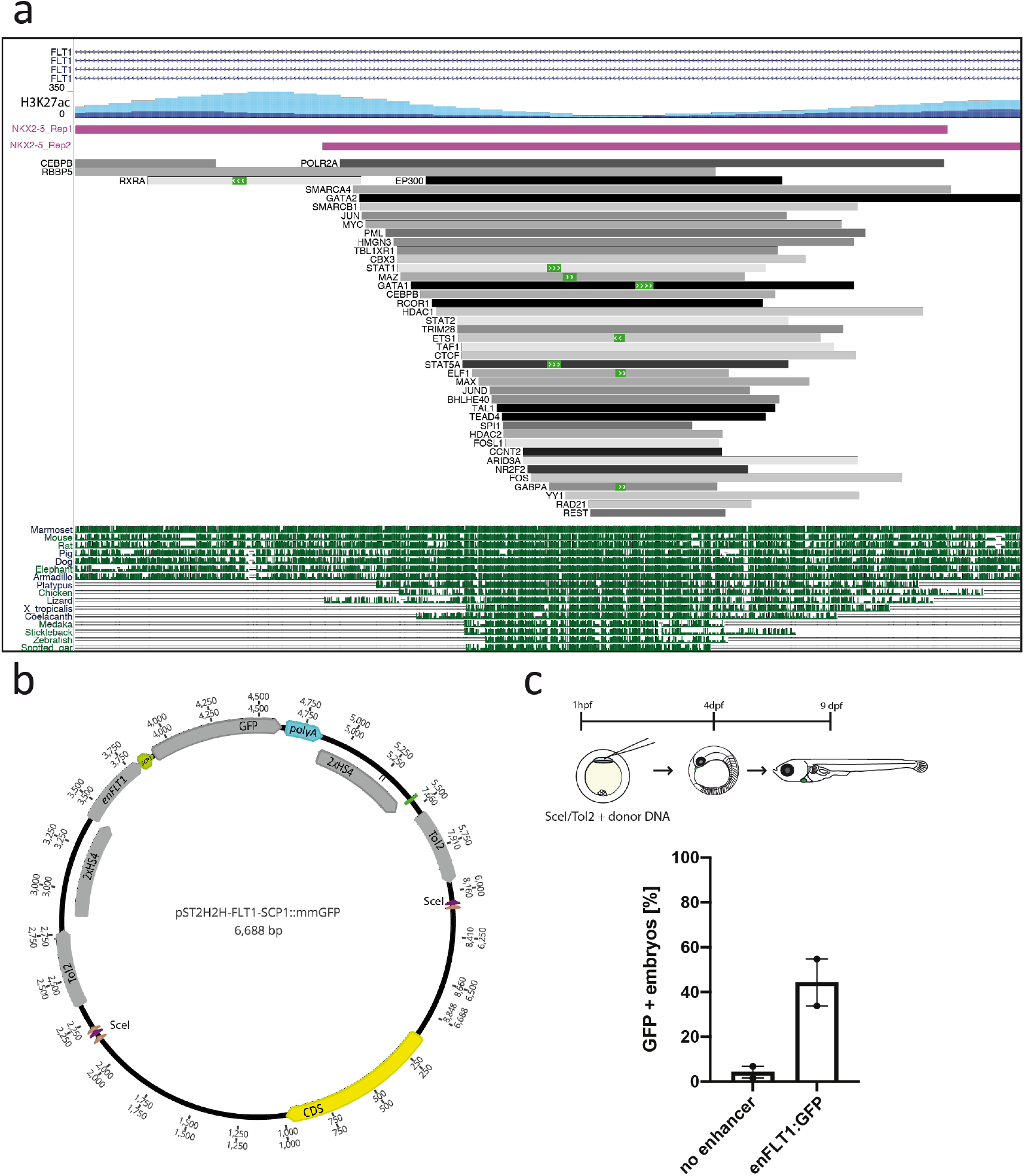
In vivo validation of *enFLT1*. a) Scheme of Transcription factor binding to *enFLT1*. Transcription factor binding overlaps with highly a conserved sequence across species and a valley between two H3K27ac activity marks suggesting open chromatin. b) Transgenesis assay plasmid containing *enFLT1*. Control vector had no RE sequence inserted. c) Plasmids were in injected in one cell stage medaka embryos. Occurrence of GFP expression was monitored until hatching. Embryos injected with the plasmid containing *enFLT1* showed higher number of GFP positive embryos compared to injection with a control vector without a putative RE.

**Figure S2:**
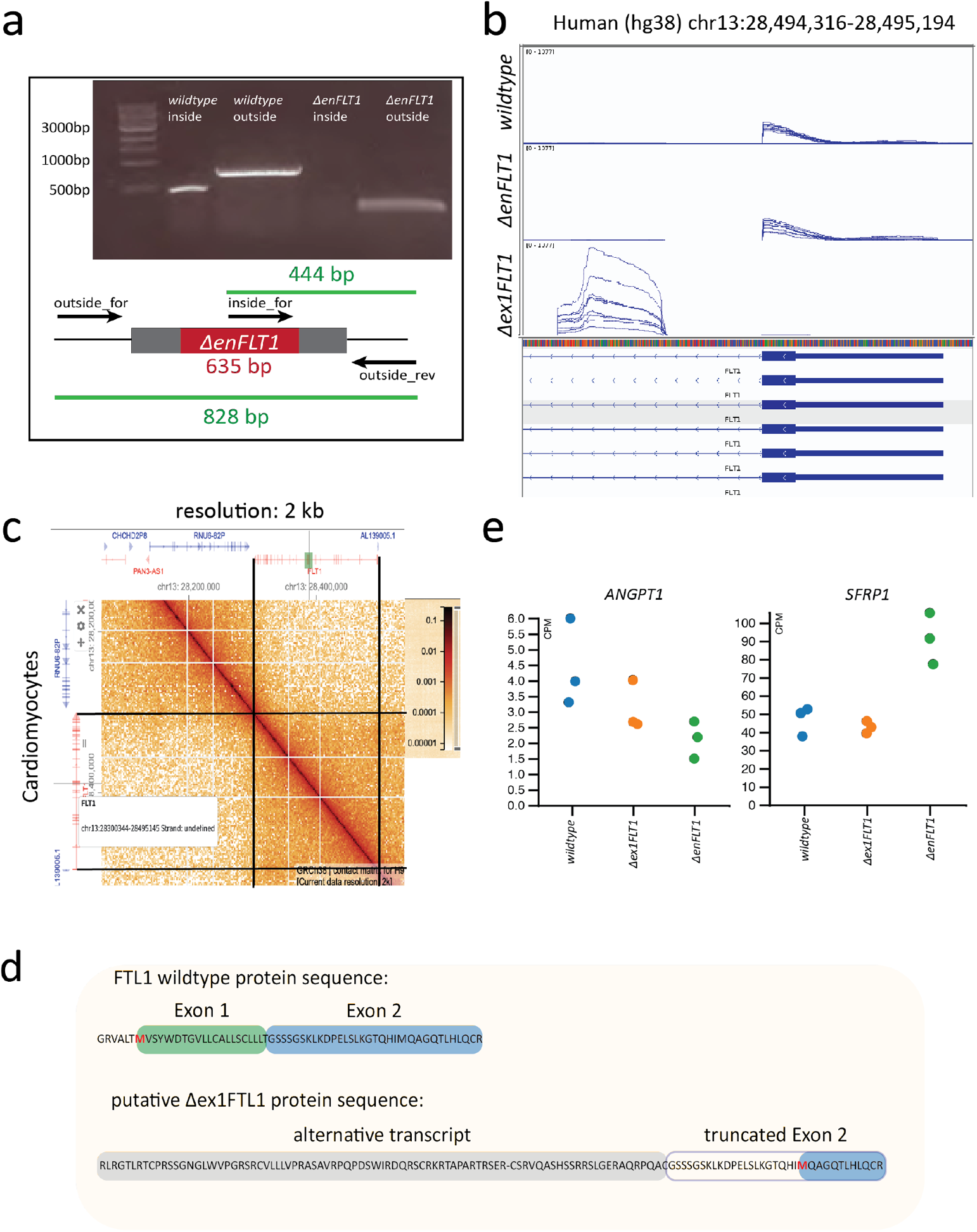
Generation of *ΔenFLT1* and *Δex1FLT1* human embryonic stem cells. a) Screening PCR targeting the deleted RE in hESCs. 3 Primer PCR approach was used to amplify gene edited locus. Band in *ΔenFLT1* outside, was cut out and sent for sequencing. b) overview of sequencing reads of exon 1 of *FLT1* in wildtype, *ΔenFLT1* and *Δex1FLT1* endothelial cells. Expression of exon 1 has been disrupted in *Δex1FLT1* and transcription starts at an alternative start site. c) HiC interaction map of targets interacting with *enFLT1* in cardiomyocytes at 2kb resolution. d) Predicted protein sequence of *FLT1* in wildtype and *Δex1FLT1*. Alternative Start codon in Exon 2 might lead to a truncated protein in *Δex1FLT1*. e) ANGPT1 and SFRP1 were identified from a list of 33 DEGs unique in *ΔenFLT1* involved in blood vessel morphogenesis.

**Figure S3:**
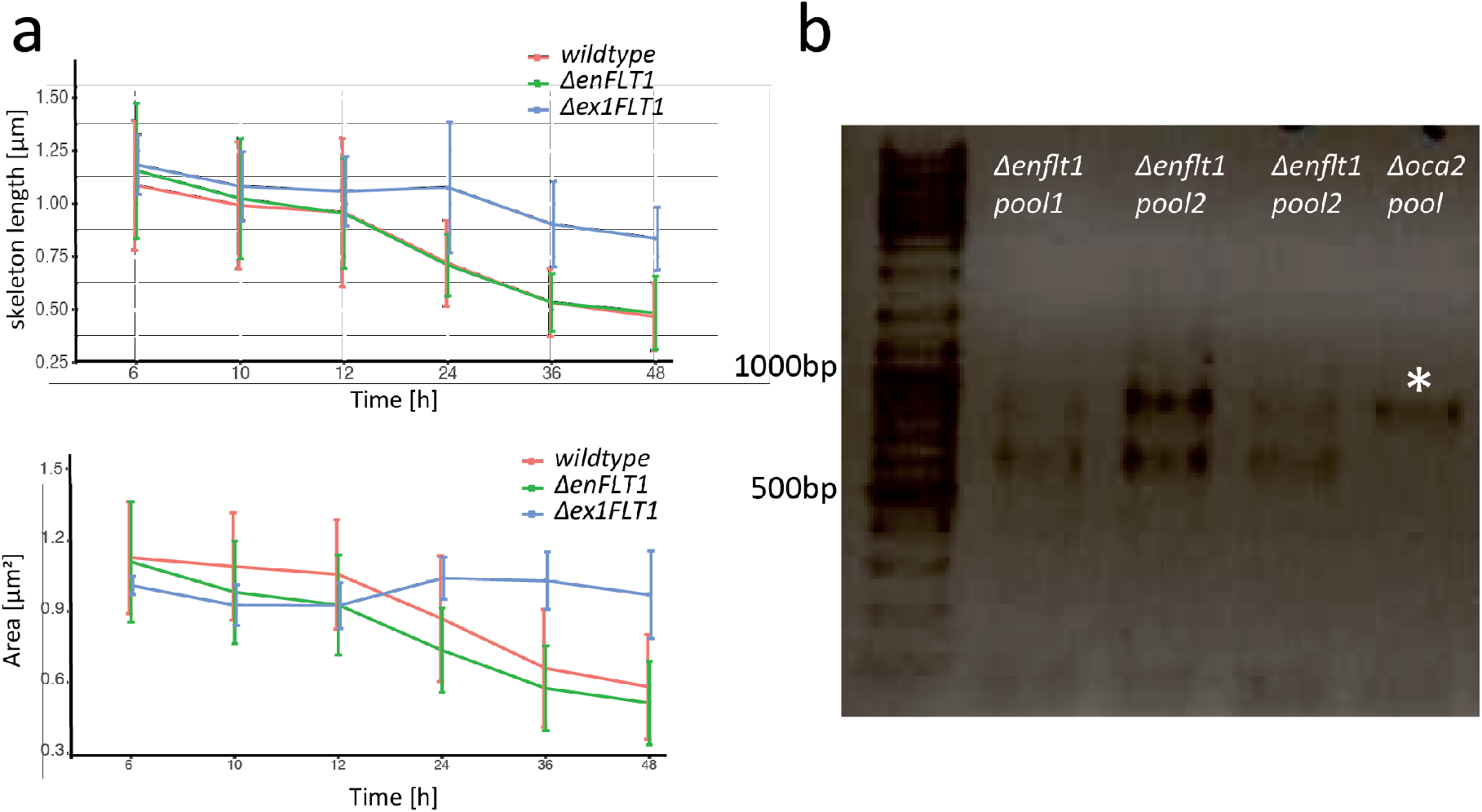
*in vitro* and *in vivo* analysis of *enFLT1* deletion. a) Assessment of normalised skeleton length and area of vessels in angiogenesis assays. Length and area are significantly different in *Δex1FLT1* compared to cell lines at 48h. b) Screening PCR of gene edited embryos injected with CRISPR/Cas9 targeting the *enflt1* or *oca2*. Asterisk indicate the wildtype band at 879bp. Excision of enhancer led to a band with a size of 634 bp. Samples contained pooled gDNA from several injected embryos.

**Table S1:**
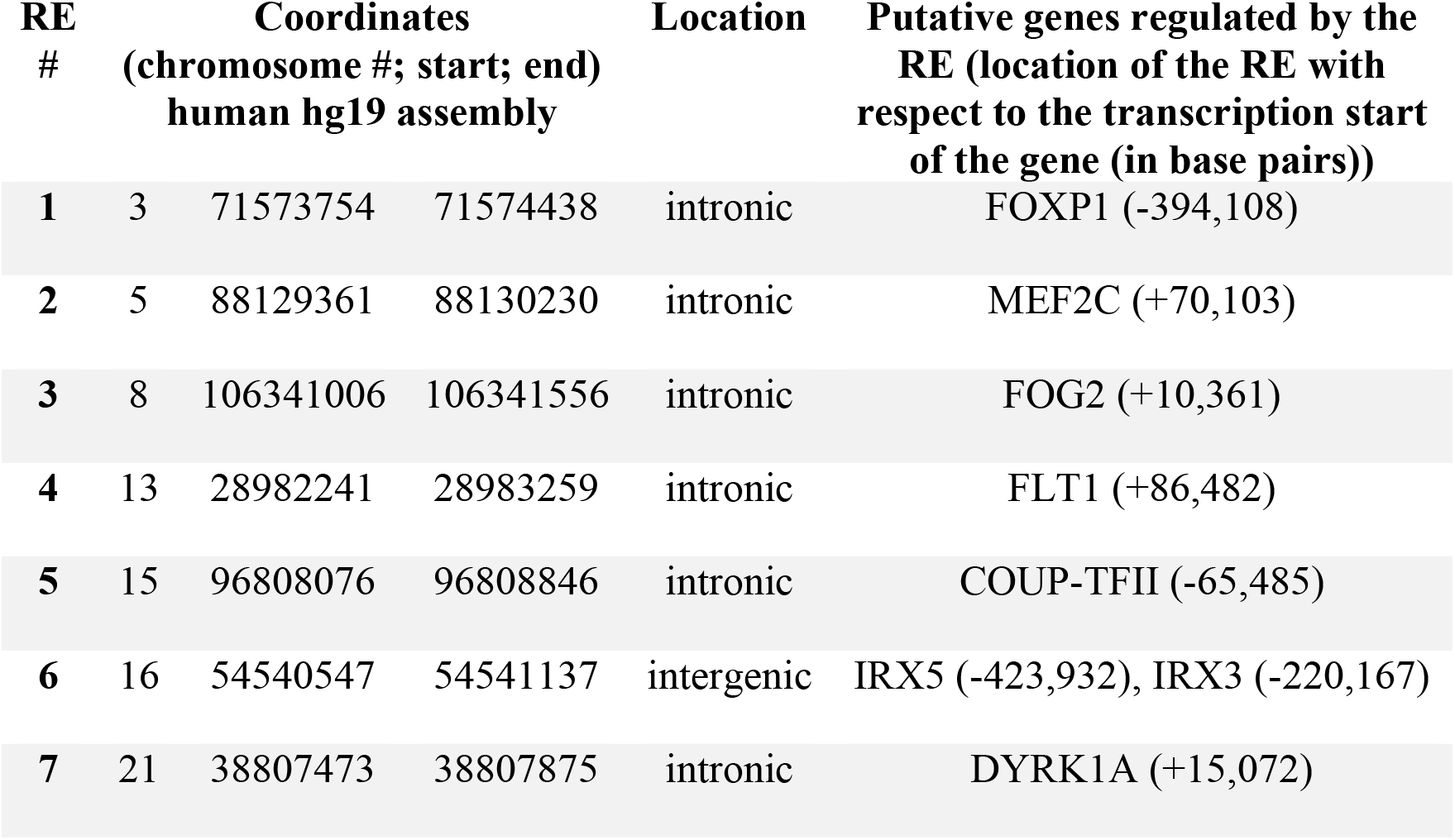
List of putative regulatory elements identified via bioinformatic mining.

